# Functional connectivity in the social perception pathway at birth is linked with attention to faces at 4 months

**DOI:** 10.1101/2025.02.27.640108

**Authors:** Katarzyna Chawarska, Angelina Vernetti, Huili Sun, Michelle Hampson, Chenhao Li, Susanne Macari, Kelly Powell, R. Todd Constable, Joseph Chang, Laura Ment, Dustin Scheinost

**Affiliations:** Child Study Center, Yale University School of Medicine, New Haven, CT; Department of Statistics and Data Science, Yale University, New Haven, CT; Department of Pediatrics, Yale University School of Medicine, New Haven CT; Department of Radiology and Biomedical Engineering, Yale University School of Medicine, New Haven

**Keywords:** neonates, faces, functional connectivity, autism, dorsal attention pathway, social perception pathway

## Abstract

**Background:** Processing faces and speech is supported by the right-lateralized social visual perception pathway (social pathway) involving medial temporal/visual 5 area (MT/V5) and superior temporal sulcus (STS). Little is known about development of the social pathway and its links with later social outcomes. We examined intrinsic functional connectivity (iFC) in the right social pathway in neurotypical neonates, compared it with iFC in the left social and bilateral dorsal attention pathways, and interrogated prospective links between iFC and social attention in neurodiverse neonates.

**Methods:** iFC in the social and dorsal pathways was measured in 517 full-term neonates from the developing Human Connectome Project (dHCP) and 73 full-term Yale neonates [M_age_=41.5 weeks (SD=1.9)]. Social attention was assessed in 36 Yale neonates at M_age_=4.2 months (SD=0.4) months.

**Results:** In the dHCP sample, the iFC indices were positive in all the pathways (all p-values < 0.001) and did not vary by sex [all p>0.275], but the iFC in the right-lateralized social pathway was higher than in the remaining pathways [all p<0.001] and was positively associated with age at scan [r(517)=0.251, p<0.011]. In the prospective neurodiverse Yale sample, the iFC in the right social pathway was positively associated with social attention at 4 months [p=0.007], and greater social attention at 4 months predicted better social functioning in the second year [p=0.010].

**Conclusions:** The early development of intrinsic functional connectivity in the right social perception pathway represents an area of interest for identifying neural mechanisms underlying emergence of atypical social attention associated with autism.

## INTRODUCTION

Dynamic human faces and speech are prioritized for processing in the attentional system shortly after birth. Neonates prefer to look at face-like stimuli compared to carefully controlled alternatives,^1^ orient more readily to child-directed over adult-directed speech,^2^ and recognize maternal voice shortly after birth.^3^ Neonates can also form intermodal pairings of faces and voices, and recognize familiar faces only when exposed to faces that speak during the familiarization.^4, 5^ Newborns prefer faces with direct gaze compared to faces with averted gaze.^6, 7^ They also integrate gaze and speech information and recognize more readily faces, which during familiarization speak *and* look at them directly compared to faces that speak while looking away.^8^ These elementary social perception skills in neonates lay the foundation for future development of face processing and social cognition in infancy.

Due to the methodological constrains related to studying task-based brain activation in the first postnatal weeks, neural mechanisms underlying early processing of social stimuli are not well understood. It has been hypothesized that the early perceptual biases for face-versus non-face patterns are mediated by subcortical structures including superior colliculus and pulvinar which are sensitive to salient features of faces including high-contrast and upper versus lower asymmetry.^9, 10^ However, it is unlikely that the range of skills exhibited by neonates can be accounted for entirely by subcortical mechanisms.^1^ Task-based functional brain imaging studies in infants suggest that 2-to 9-month-old infants activate face selective areas including fusiform gyrus in response to unimodal static faces.^11, 12^ It remains unknown, however, to what extent the cortical areas known to respond preferentially to faces and speech are functional in neonates.

In the human brain, processing of dynamic social information is supported by a social perception pathway that extends from the visual areas through motion sensitive medial temporal (MT/V5) and superior temporal sulcus (STS) areas.^13^ While face and speech specific responses are already seen at the MT/V5 level,^14^ the STS specializes in dynamic aspects of social perception including biological motion, dynamic faces, and speech, and their multimodal combinations.^14, 15^ The right-lateralized posterior STS (pSTS) is involved in face processing^14, 16^ and biological motion,^14, 17^ the medial STS (mSTS) responds to speech stimuli and faces,^14^ and the anterior STS (aSTS) has been implicated in gaze direction perception^18^ and speech.^14^ The STS is characterized by a posterior to anterior gradient with more basic perceptual information represented in the posterior parts of the pathway to more abstract or conceptual representation of social stimuli moving from MT/V5 toward aSTS.^13, 19^

The STS has been an area of interest in autism for several decades. Children with autism display atypical developmental trajectories of resting state connectivity in the STS.^20^ Alterations in resting-state functional connectivity between STS and other brain areas specializing in social perception and cognition including the frontoparietal areas,^21^ inferior parietal lobule and premotor areas,^22^ the amygdala, ventromedial prefrontal cortex, and fusiform gyrus^22–26^ have been reported extensively in older children and adults with autism. Moreover, individuals with autism display hypoactivation in the voice selective regions of the right and left STS.^27^ To the best of our knowledge, only one study examined resting-state connectivity *within* the social pathway in autism. Using the ABIDE sample, Li and colleagues demonstrated that children with autism demonstrate decreased connectivity between the right pSTS and mSTS compared to typically developing controls.^28^ The clinical significance of this finding is highlighted by the association between the lower pSTS-mSTS connectivity and higher severity of autism symptoms in the social domain.

In the preset study, we contrast the right-lateralized social pathway with the pathway involving left STS, which is concerned with highly social but largely auditory information involving speech and multimodal integration of speech and sematic concepts^29–32^ as well as gaze cues.^33^ This contrast is essential, as it is not clear to what extent the functional differentiation between the left and right STS is present in neonates. We also contrast the social pathway with the dorsal attention pathway. The dorsal pathway extends from MT/V5 and consists of the inferior parietal sulcus (IPS) and frontal eye fields (FEF),^34, 35^ is activated during tasks involving visual-spatial top-down attention but also feature- and object-based attention, encodes and maintains preparatory signals and modulates top-down visual and other sensory regions.^34, 36–38^ The dorsal pathway is bilateral,^37^ though spatial attention signals are largely carried in the right pathway.^39^ Although the DAN is not specifically involved in processing social stimuli, it has been implicated in autism both on the functional connectivity^40^ and behavioral ^41, 42^ levels.

The present study examined for the first time intrinsic functional connectivity in the social visual pathways in a large sample of typically developing neonates. Considering the potential relevance of the social pathway for processing multimodal social cues observed in newborns,^4–8^ we hypothesized that the nodes within the right-lateralized social pathway will show positive coactivation, that the connectivity within the social pathway will increase with age as a function of exposure to a highly dense social environment,^43^ and will be stronger than in the control pathways. Lastly, leveraging a prospective neurodiverse sample of neonates, we examined the links between functional connectivity in the social pathway and attention to dynamic multimodal faces at 4 months. Selective attention to dynamic multimodal faces is already critically diminished in autism in presymptomatic infants and in symptomatic toddlers.^44–46^ We hypothesized that if the strength of connectivity within the right social pathway at birth facilitates development of social perception, there will be a significant positive association with later social attention. Given that social pathway is thought to be right lateralized with regard to processing dynamic faces and speech,^13^ we hypothesized that these correlations will not be observed in control pathways.

## METHODS AND MATERIALS

### Participants

#### Developing human connectome project (dHCP) Neonatal Sample

The dHCP is an observational, cross-sectional Open Science program approved by the UK National Research Ethics Authority (**Table 1)**. The dHCP sample consisted of 517 full-term neonates (54% male) with PMA at birth ranging from 37.0 to 42.7 weeks Infants were recruited from the postnatal wards and scanned at 37.4 to 44.9 weeks PMA. The dHCP sample exclusion criteria include a history of severe compromise at birth requiring prolonged resuscitation, a diagnosed chromosomal abnormality, or any contraindication to MRI scanning.

**Table 1.** Sample Characteristics.

|  | dHCP Sample | Yale Sample: NN Scan | Yale Sample: 4-months eye-tracking data |
| --- | --- | --- | --- |
| N | 517 | 73 | 36 |
| Female (N, %) |  |  |  |
| Male (N, %) | 281 (54%) | 40 (55%) | 18 (50%) |
| PMA at Birth | 39.95 (1.27) | 39.74 (0.86) | 39.88 (1.00) |
| PMA at Scan | 41.28 (1.73) | 43.92 (1.32) | 44.13 (1.21) |
| FtF Motion | -- | 0.03 (0.02) | 0.03 (0.02) |
| Minutes Rest | -- | 11.44 (1.99) | 11.35 (2.27) |
| Family Hx of Autism | -- | 29% (21/73) | 28% (10/36) |
| Developmental outcomes |  |  |  |
| Typical | -- | 53% (39/73) | 56% (20/36) |
| Atypical* | -- | 32% (23/73) | 28% (10/36) |
| Unknown** | -- | 15% (11/73) | 17% (6/36) |
| Follow-Up Visit Age (months) | -- | 22.98 (4.90) | 22.49 (5.67) |
| MSEL Verbal DQ | -- | 100.64 (22.90) | 98.65 (22.48) |
| MSEL Nonverbal DQ | -- | 102.11 (13.17) | 104.44 (15.07) |
| ADOS-2 Total raw score | -- | 4.08 (3.07) | 3.69 (2.63) |
| VABS Comm SS | -- | 97.53 (13.85) | 97.91 (15.63) |
| VABS DLS SS | -- | 93.54 (14.99) | 95.60 (15.59) |
| VABS Soc SS | -- | 96.54 (12.56) | 97.97 (12.76) |
| FYI Social/Comm Score | -- | 5.10 (6.58) | 4.85 (5.89) |
| FYI Regulatory Score | -- | 5.00 (7.37) | 4.87 (6.02) |

\* Atypical group includes 3 children with autism in the full sample and 2 children with autism in the follow-up sample
\*\*Lost to follow-up

#### Yale Neonatal Sample

Seventy-seven healthy infants were scanned between April 2018 and July 2022. Four infants awoke before adequate functional data could be collected; thus, usable data were available for 73 (55% males) of neonates (**Table 1**). Inclusion criteria were singleton pregnancy, term birth (born after 37 weeks gestation), and appropriate for gestational age. Exclusion criteria were: 1) congenital infections; 2) non-febrile seizure disorder; 3) hearing loss; 4) visual impairment; 5) the presence of known chromosomal abnormality; 6) prenatal exposure to illicit drugs; 7) major psychotic disorder in first degree relative; and 8) contraindications to MRI including non-removable metal medical implants. PMA at birth ranged from 37.6 to 41.9 weeks and the scans were conducted between 40.7 and 46.4 postmenstrual weeks Out of 73, 21 (29%) of neonates had a family history of autism in the first-or second-degree relatives. Due to genetic factors, infants with familial history of autism have an elevated likelihood of developing autism or related challenges referred to as broader autism phenotype. Amongst children with first degree relatives with autism (e.g., parent or sibling), approximately 20% are likely to be diagnosed with the condition as well and 30-40% to exhibit BAP features.^47, 48^ At follow-up in the second year of life, 23 out of 72 participants (32%) evidenced some developmental vulnerabilities ranging from autism (n=3) to global or specific developmental delays (n=20) (Table 1). Out of 73 neonates, 36 attended the 4-month visit and contributed valid data to the SSA 4.0 procedure. Of the 37 who did not contribute eye-tracking data, 15 skipped the visit or did not participate in the SSA 4.0 procedure due to technical reasons, and 22 did not contribute valid eye-tracking data due to arousal regulation issues or inattention.

### Imaging Acquisition

#### dHCP Cohort

Imaging was acquired at the Evelina Newborn Imaging Centre, Evelina London Children’s Hospital, using a 3 T Philips Achieva system (Philips Medical Systems). All infants were scanned without sedation in a scanner environment, including a dedicated transport system, positioning device, and a customized 32-channel receive coil with a custom-made acoustic hood. MRI-compatible ear putty and earmuffs were used to provide additional acoustic noise attenuation, and infants were fed, swaddled, and positioned in a vacuum jack prior to scanning to provide natural sleep.^49^ High temporal resolution multiband EPI (TE=38 ms; TR=392 ms; MB factor=9x; 2.15 mm isotropic) specifically developed for neonates was acquired for 15 min.

#### Yale Neonatal Project

Participants were scanned without sedation during natural sleep using the feed-and-wrap protocol.^50^ Infants were fed, bundled with multiple levels of ear protection, and immobilized in an MRI-safe vacuum swaddle. Heart rate and O_2_ saturation were continuously monitored during all scans. The scans were performed using a 3 Tesla Siemens (Erlangen, Germany) Prisma MR system with a 32-channel parallel receiver head coil. Functional runs were acquired using a multiband T2*-sensitive gradient-recalled, single-shot echo-planar imaging pulse sequence (TR = 1 s, TE = 31 ms, FoV = 185 mm, flip angle 62°, multiband = 4, matrix size 92 × 92). Each volume consisted of 60 slices parallel to the bi-commissural plane (slice thickness 2 mm, no gap). We collected 4-5 functional runs, each comprised of 360 volumes. The mean frame-to-frame displacement was calculated for each run for every individual and runs with a mean frame-to-frame displacement greater than 0.10 mm were removed from further analysis. Each neonate had on average 11.4 min (SD = 2.0) of usable functional data with an average frame-to-frame displacement of 0.03 (SD=0.02, Min: 0.01, Max: 0.10). High-resolution T1-and T2-weighted 3D anatomical scans were acquired using an MPRAGE sequence (TR=2400 ms, TE=1.18 ms, flip angle=8°, thickness=1 mm, in-plane resolution=1 mm x 1 mm, matrix size=256 × 256) and a SPACE sequence (TR=3200 ms, TE=449 ms, thickness=1 mm, in-plane resolution=1 mm x 1 mm, matrix size=256 × 256).

### Image Processing

For the dHCP data, the dHCP functional pipeline was used to align the functional images to the dHCP template space^53^. For the Yale data, functional images were aligned to a custom infant template using a series of linear and nonlinear registrations. These registrations were calculated independently and combined into a single transform. This allows the participant images to be transformed to common space with only one transformation, reducing interpolation error. Functional images were linearly registered to the anatomical image, which was nonlinearly registered to the infant template using a previously validated algorithm.^54^ Similarly, the same algorithm registered the infant templates for the dHCP and Yale datasets to the MNI brain.

#### Connectivity processing

The dHCP data was preprocessed with the dHCP functional pipeline^53^, including echo distortion correction, motion correction, independent component analysis (ICA) denoising, and registered to individual T2w native space. The Yale data were processed using a previously validated pipeline.^52^ Functional images were slice-time and motion corrected using SPM8.

Next, images were iteratively smoothed until the smoothness of any image had a full-width half maximum of approximately 6 mm using AFNI’s 3dBlurToFWHM. This iterative smoothing reduces motion-related confounds.^55^ For both datasets, all further analyses were performed using BioImage Suite^56^ unless otherwise specified. Several covariates of no interest were regressed from the data including linear and quadratic drifts, mean cerebral-spinal-fluid (CSF) signal, mean white-matter signal, and mean-gray matter signal. For additional control of possible motion-related confounds, a 24-parameter motion model (including six rigid-body motion parameters, six temporal derivatives, and these terms squared) was regressed from the data. The data were temporally smoothed with a Gaussian filter (approximate cutoff frequency=0.12Hz). A canonical gray matter mask defined in common space was applied to the data, so only voxels in the gray matter were used in further calculations.

#### Functional connectivity

The coordinates of the nodes in the social and dorsal networks was drawn from Lanhakoski et al ^17^ (see **Figure 2** for details). After the seeds for each of the three pathways were warped from MNI space into a single participant’s space, the time course for each seed region was then computed as the average time course across all voxels in the reference region. The time courses were correlated between seed pairs and transformed to z-values using Fisher’s transform. Following Li et al.,^41^ intra-network connectivity strength for each pathway was defined as the average of the individual connections within each pathway, resulting in four functional connections per individual. For the social pathway, we examined connectivity between MT/V5 and pSTS, between pSTS and mSTS, and between mSTS and aSTS in both hemispheres to investigate how neonates process and integrate information along the pathway. The same strategy was applied for the dorsal pathway in both hemispheres (MT/V5 to IPS and IPS to FEF).

**Figure 1.**
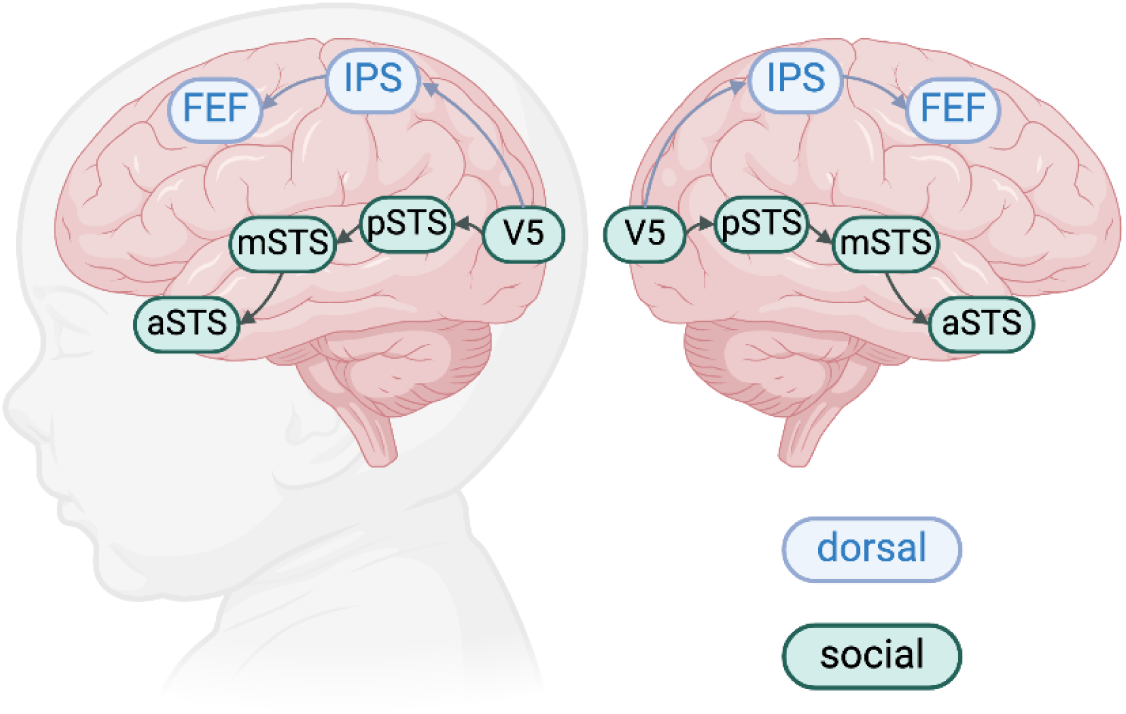
Schematic representation of the social perception and dorsal attention pathways in the brain. The social perception pathway extends from the motion sensitive medial temporal/visual 5 (MT/V5) area through the posterior, medial, and anterior superior temporal sulcus (STS). The dorsal attention pathway extends from the MT-V5 through the interparietal sulcus (ISP) into the frontal eye fields (FEF).

**Figure 2.**
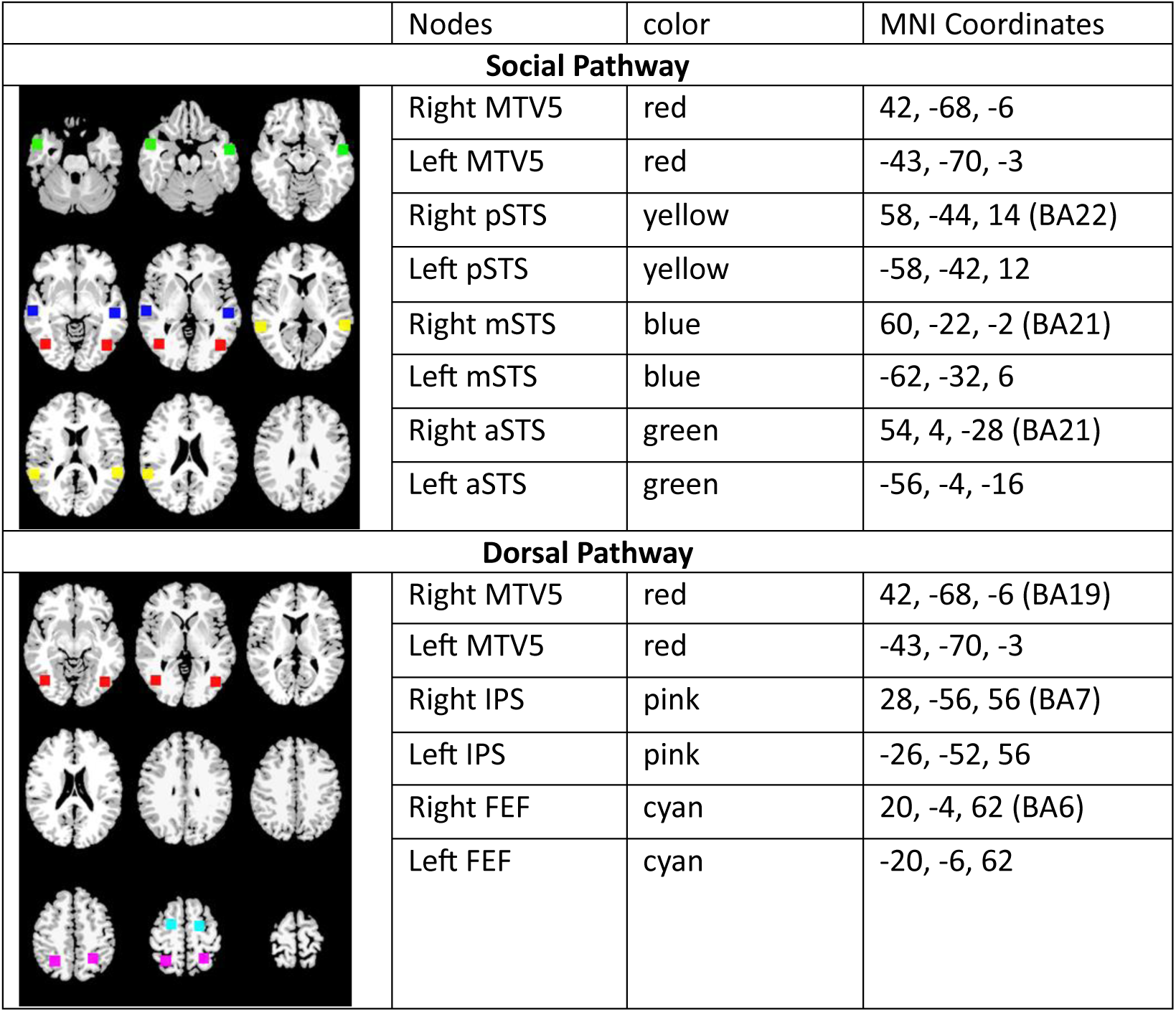
The MNI coordinates for the social perception and dorsal attention pathways.

### Social Attention Procedure at 4 months

At 4 months infants were administered the selective social attention task, version 4.0 (SSA 4.0). The infants completed the eye-tracking procedure at 4.2 months (SD=0.4). Gaze location was captured using the SR EyeLink 1000 Plus Hz (SR Research Ltd) eye tracker using a 5-point calibration procedure.

The stimuli consisted of 8-second videos of a person surrounded by 4 distractor toys (**Figure 3**). Sessions were conducted in a dark, sound-proof room with a 24” widescreen LCD monitor. Infants sat in a high-chair approximately 60 centimeters from the screen. Each session began with a short cartoon video followed by a calibration procedure. Each infant was presented with 4 8-second trials of a stimulus where a person spoke using child-directed speech while looking directly at the camera. The trials were separated by 500 millisecond breaks consisting of a black screen with no fixation cross or attention-getters. The primary dependent measure was the proportion of looking at a person’s face while speaking and looking directly at the camera (%Face) where duration of looking at the face was standardized by the total duration of looking at the scene.

**Figure 3.** A screen shot depicting the experimental layout for the Selective Social Attention task implemented at 4 months.

**Figure 4.**
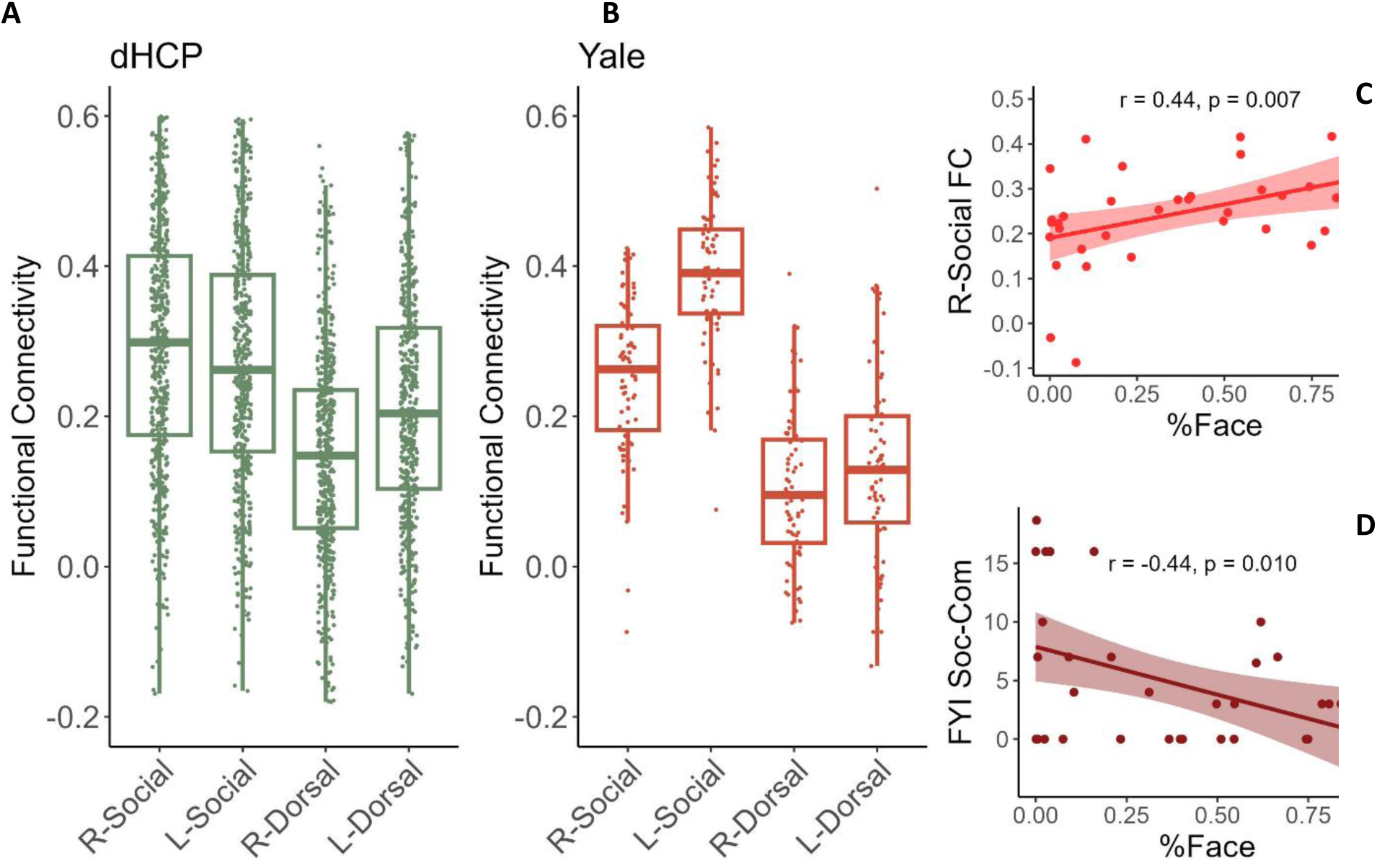
Boxplots representing resting-state functional connectivity in the social perception and dorsal attention pathways in neonates drawn from the Developing Human Connectome Project (dHCP) (Panel A in green) and Yale Neonatal Project (Panel B in red). Plots represeting Pearson r correlations between proportion of looking at the face at 4 months (%Face) and functional connectivity in the right social pathway (Panel C) and Fist Year Inventory social communication scores in the second year of life (Panel D).

### Assessment of Behaviors Associated with Autism in the Second Year of Life

Behaviors that are associated with autism in toddlers were assessed using the First Year Inventory 2.0 (FYI), a parent questionnaire consisting of the Social Communication (28 items) and Sensory-Regulatory (24 items) scales.^55, 56^ The FYI measure is sensitive to autism-specific behaviors in the general population and amongst infant with family history of autism,^57–61^ and shows positive correlations with concurrent measures of autism symptom severity based on the ADOS-2 Toddler Module^62^ or the Autism Observation Scale for Infants (AOSI).^63^ Unlike the ADOS-2 and AOSI, the FYI was designed to quantify autism traits in the general population and thus, is an appropriate measure for neurodiverse samples. Risk scores were generated based on assigned risk points derived from the normative sample.^55^ Each scale scores range from 0 to 50 on a semi-logarithmic scale, with higher scores indicating more symptoms.

### Statistical Analysis

To examine strength of connectivity within each pathway, we performed a series of t-tests to compare the iFC values to zero. Planned paired t-tests compared iFC in the right social pathway with the remaining four pathways. Sex effects on iFC were examined using a series of general linear models.

Exploratory analyses examining links between social pathway iFC and behavioral outcomes were performed using Pearson’s r correlation analysis. Age at scan (PMA) and frame-to-frame displacement were included into the analyses as indicated. Only the effects that survive the Bonferroni correction for multiple comparisons are interpreted.

### Data and code availability

The YaleACE neonatal datasets will be released on https://nda.nih.gov/ following an embargo period. The dHCP data can be accessed at http://www.developingconnectome.org/project/. The image analysis software (BioImage Suite) can be found at https://medicine.yale.edu/bioimaging/suite/ and https://bioimagesuiteweb.github.io/webapp/index.html.

## RESULTS

### Developing Human Connectome Project Neonatal Sample

#### Intrinsic functional connectivity

**Table 2** presents the descriptive statistics for social and dorsal pathways in the dHCP sample. The average iFC values were significantly greater than 0 in the R-social [t(1, 516) = 39.18, p<0.001], L-social [t(1, 516) = 35.51, p<0.001], L-dorsal, [t(1, 516) = 36.51, p<0.001], and R-dorsal, [t(1, 516) = 22.40, p<0.001] pathways. All the effects remained statistically significant after Bonferroni correction for multiple comparisons (p<.013).

**Table 2.** Mean (SD) functional within network connectivity the right and left social perception and dorsal attention pathways in the dHCP and Yale samples.

| Variable | dHCP Sample |  |  | Yale Sample |  |  |
| --- | --- | --- | --- | --- | --- | --- |
|  | N | M(SD) | H0=0<br>p-value | N | Mean | H0=0<br>p-value |
| R-Social | 517 | 0.30 (0.17) | <0.001 | 73 | 0.25 (0.10) | <0.001 |
| L-Social | 517 | 0.27 (0.17) | <0.001 | 73 | 0.39 (0.10) | <0.001 |
| R-Dorsal | 517 | 0.15 (0.15) | <0.001 | 73 | 0.10 (0.11) | <0.001 |
| L-Dorsal | 517 | 0.21 (0.16) | <0.001 | 73 | 0.13 (0.13) | <0.001 |

Planned pairwise comparisons indicated that R-social iFC was greater than L-social [M_diff_ = 0.027 (SD=0.182), t(1, 516) = 3.34), p<0.001], R-dorsal [M_diff_ = 0.154 (SD=0.206), t(1, 517) = 17.00, p<0.001], and L-dorsal [M_diff_ = 0.086 (SD=0.219), t(1, 516) = 8.93, p<0.001] iFC. All the effects remained statistically significant after Bonferroni correction for multiple comparisons (p<.017).

For statistics describing individual edges, please see **Table S1** in the **Supplement**.

#### Sex differences

A series of general linear model analyses controlling for the effects of PMA at scan, indicated no significant effects of sex for the any of the five pathways with all p-values greater than 0.275 (**Table 3**).

**Table 3.** A comparison of the iFC in the social and dorsal pathways in male and female neonates while controlling for post-menstrual age (PMA) at scan in the Developing Human Connectome Project sample (top) and the ACE sample (bottom).

| **dHCP Sample** |  |  |  |  |  |
| --- | --- | --- | --- | --- | --- |
|  | Female |  | Male |  |  |
| Visual Pathway | N | M (SD) | N | M (SD) | p-value |
| R-Social | 236 | 0.30 (0.18) | 281 | 0.30 (0.17) | 0.799 |
| L-Social | 236 | 0.27 (0.17) | 281 | 0.28 (0.17) | 0.275 |
| R-Dorsal | 236 | 0.14 (0.15) | 281 | 0.15 (0.15) | 0.486 |
| L-Dorsal | 236 | 0.21 (0.17) | 281 | 0.22 (0.16) | 0.182 |
| **Yale Sample** |  |  |  |  |  |
| Visual Pathway | N | Mean | N | Mean | p-value |
| R-Social | 33 | 0.26 (0.11) | 40 | 0.24 (0.10) | 0.493 |
| L-Social | 33 | 0.39 (0.10) | 40 | 0.39 (0.10) | 0.956 |
| R-Dorsal | 33 | 0.11 (0.09) | 40 | 0.09 (0.13) | 0.369 |
| L-Dorsal | 33 | 0.12 (0.13) | 40 | 0.14 (0.13) | 0.849 |

#### Associations with age

The iFC in the R-social [r(517) = 0.251, p<0.001] and L-social [r(517) = 0.144, p=0.001] pathways showed statistically significant correlations with age at scan (**Table 4**). There were no significant correlations between the L-dorsal [r(517) = 0.075, p=0.087 and R-dorsal [r(517) = 0.007, p=0.867] iFC with age. The correlations observed in the left and right social pathways remained significant after Bonferroni correction for multiple comparisons (p<.017).

**Table 4.** Pearson’s r correlation coefficient examining associations between post-menstrual age (PMA) at the time of the scan with the intrinsic functional connectivity in the social perception and dorsal attention pathways in the dHCP (N=517) and the ACE (N=73) samples.

| Visual Pathway | Statistic | dHCP sample (N=517) | Yale sample (N=73) |
| --- | --- | --- | --- |
|  |  | PMA at Scan | PMA at Scan |
| R-social | $r$ | 0.251 | 0.203 |
| | $p$ | **<0.001** | 0.085 |
| L-social | $r$ | 0.144 | -0.017 |
| | $p$ | **0.001** | 0.885 |
| R-dorsal | $r$ | 0.007 | 0.017 |
| | $p$ | 0.867 | 0.890 |
| L-dorsal | $r$ | 0.075 | 0.036 |
| | $p$ | 0.088 | 0.765 |

### Yale Neonatal Sample

#### Intrinsic functional connectivity, associations with age, and sex effects

Similarly, as in the dHCP sample, in the ACE neonates, there was robust evidence of functional coactivation between the nodes of the R-social [t(72) = 20.68, p<0.001], L-social [t(72) = 34.35, p<0.001], R-dorsal [t(72) = 7.39, p<.0.001], and L-dorsal [t(72) = 8.90, p<0.001] pathways (**Table 2**). As in the dHCP sample, there were no significant effects of sex on iFC in any of the pathways (**Table 3**). However, unlike in the Lastly, in the smaller sample, associations with age were largely not statistically significant, though the positive association between R-social pathway and PMA showed similar magnitude of the effect as in the dHCP sample and a trend toward statistical significance (**Table 4**).

Pairwise comparisons between the right social pathway and the dorsal left [M_diff_ = 0.119, (SD=0.164), t(1, 72) = 6.20), p<0.001] and right dorsal [M_diff_ = 0.154 (SD=0.136), t(1, 72) = 9.71), p<0.001] pathways indicated that the connectivity in the right social pathway was higher than in the dorsal pathways. However, in this neurodiverse sample, connectivity in the left social pathway was higher than in the right social pathway [M_diff_ = 0.143 (SD=0.121), t(1, 72) = 10/05), p<0.001].

#### Associations between social pathway and social attention at 4 months

Out of 73 neonates, 36 attended the 4-month visit and contributed valid data to the SSA 4.0 procedure. The retained infants did not differ from not-retained infants in terms of PMA at scan [p=0.179] or iFC in the R-social [p=0.47], L-social [p=0.226], R-dorsal [p=0.097], and L-dorsal [p=0.294], pathways at birth. They also did not differ in the verbal [p=0.471] and nonverbal [p=0.135] DQ or the FYI social communication [p=0.762] or FYI emotion regulation [p=0.893] scores at outcome.

Overall, the 4-month-old infants attended to the dynamic speaking faces 37% (SD=32) of the time they looked at the screen. After controlling for PMA and frame-to-frame displacement, greater iFC in the R-social pathway in was positively associated with attention to dynamic faces at 4 months, r(36) = 0.454, p=0.007 (**Figure 3**) and the effect remained significant after Bonferroni correction for multiple comparisons (p<.013). The clinical significance of this finding is underscored by the association between %Face at 4 months and later FYI social communication score (r(33) = −0.441, p=0.010) but not with the FYI sensory regulation score (r(31) = −0.003, p=0.987. There were no significant associations between the L-social pathway [r(36) = 0.104, p=0.557], L-dorsal [r(36) = −0.007, p=0.969], or R-dorsal [r(36) = - 0136, p=0.429] pathways with social attention at 4 months.

## DISCUSSION

The study examined the social perception pathway specializing in processing dynamic faces, speech, and gaze stimuli in neonates. The pathway extends from the right visual and motion sensitive MT/V5 area through the posterior, medial, and anterior STS.^13^ We report for the first time that in typically developing neonates the right-lateralized social pathway is functional shortly after birth and has higher functional connectivity than the left social and dorsal attention pathways. Its connectivity increases rapidly with postnatal age and does not vary by sex. Similar trends related to overall strength of connectivity and sex were observed in an independent sample of neurodiverse neonates, though in this sample the right social pathway was less connected than the left social pathway. Crucially, we report for the first time, that the strength in functional connectivity in the right social pathway predicts social attention at 4 months, which in turn associates with better social functioning at 20 months. Importantly, these associations were not observed in the left STS or dorsal pathways.

In the first three months of life, infants spend nearly a quarter of their waking hours looking at dynamic, large (viewed up close) multimodal faces that include both eyes.^43^ No other object category is as frequently viewed during this period as faces.^43^ Newborns demonstrate advanced perceptual skills in processing of this dense social visual and auditory input and have a capacity to integrate this input in service of learning and memory.^1, 2, 4–7^ Here we demonstrate that the cortical areas located along the social pathway specializing in processing dynamic multimodal faces show strong coactivation shortly after birth and we propose that this pathway supports social perception skills observed in neonates.

While this hypothesis awaits a direct empirical test in a context of task-based fMRI protocol, supporting evidence for the hypothesis comes from several sources. First, other cortical networks such as default mode, salience, and executive control^64^ as well as language^65^ networks evidence positive co-activation patterns during the prenatal to postnatal transition, and associate with later outcomes.^66^ Second, studies utilizing functional near infrared spectroscopy (fNIRS)^67^ and electrophysiological (EEG)^68^ methods demonstrate that the broadly outlined occipitotemporal cortices areas including superior temporal gyrus and superior temporal sulcus activate preferentially in response to faces shortly after birth. Lastly, here we demonstrate that greater functional connectivity within the right-lateralized social pathway at birth predicts better ability to attend to dynamic, multimodal faces four months later. Albeit preliminary, this study offers the first evidence of a highly relevant to survival social perception pathway in neonates and its importance for the development of social attention in infancy.

The present findings shed light not only on development of social perception in typically developing children, but also may contribute to understanding of the brain processes underlying one of the best replicated biomarkers in autism: limited attention to dynamic multimodal faces. Limited attention to dynamic faces had been replicated across cohorts, laboratories, and tasks, and has been reported in presymptomatic infants later diagnosed with autism,^44–46^ and in toddlers ^69, 70^ and school age children with autism.^71, 72^ We demonstrate for the first time that attention to dynamic multimodal faces at 4 months relates to the strength of connectivity within the right social perception pathway at birth in neurodiverse infants. The neurodiverse infants include those with family history of autism and thus with elevated likelihood for developing autistic and broader phenotype features. Although face processing skills develop into adolescence, the first months of life are highly formative for social behavior, and their importance is highlighted by the findings linking attention to faces with later social outcomes in the present sample and in other studies.^73–75^ The significance of our findings is further strengthened by a recent report that underconnectivity between the nodes in the right social perception pathway in older children with autism associate with higher autism symptom severity, and more broadly, social vulnerability trait in children with and without autism.^28^

### Limitations and future directions

Although the overall characterization of the social pathway in neonates is based on a large sample of neonates, the associations with later outcomes were assessed in a smaller sample and thus, more sophisticated outcome modeling approaches were not feasible. the patterns of connectivity strength between the left and right social perception pathways were reversed in the neurodiverse neonates compared to the neurotypical dHCP sample. The significance of this finding remains to be clarified. Interpretation of the present study will benefit greatly from a replication in a larger sample of neonates with risk factors for social vulnerabilities, which hopefully will strengthen and further clarify the link between neonatal connectome and the attentional autism biomarker.

### Conclusions

The present study demonstrates that a functional brain pathway tuned to processing multimodal social stimuli that are most common in the newborn’s environment shows is highly integrated and associates robustly with later social outcomes relevant to autism.

## Acknowledgements

None.

## Additional Contributions

We thank the children and their families for participating in the study. We acknowledge the clinical team of the Yale Social and Affective Neuroscience of Autism Program including Chelsea Morgan, Megan Lyons, and Amy Carney, for their contribution to sample characterization and data collection.

## Author Information

Authors do not have any competing financial interests to disclose.

## Correspondence

Correspondence should be addressed to

## Data Availability

Data will be available upon reasonable request to the Authors contingent upon a formal data sharing agreement, project outline, and approval from the requesting researcher’s ethics committee.

