## Supplement for "Functional connectivity in the social perception pathway at birth is linked with attention to faces at 4 months"

Table S1. Pairwise correlations between the nodes within the social perception and dorsal attention pathways in the dHCP and Yale Neonatal samples.

| Sample | | dHCP | | | Yale | | |
|---|---|---|---|---|---|---|---|
| Edge | Pathway | N | Mean | SD | N | Mean | SD |
| RMTV5 - RpSTS | Social | 517 | 0.06 | 0.19 | 73 | 0.03 | 0.17 |
| RpSTS - RmSTS | | 517 | 0.52 | 0.29 | 73 | 0.48 | 0.20 |
| RmSTS - RaSTS | | 517 | 0.32 | 0.25 | 72 | 0.24 | 0.16 |
| LMTV5 - LpSTS | | 517 | 0.06 | 0.20 | 73 | 0.08 | 0.14 |
| LpSTS - LmSTS | | 517 | 0.40 | 0.27 | 73 | 0.48 | 0.19 |
| LmSTS - LaSTS | | 517 | 0.36 | 0.25 | 73 | 0.62 | 0.16 |
| RMTV5 - RIPS | Dorsal | 517 | 0.10 | 0.21 | 73 | 0.18 | 0.16 |
| RIPS - RFEF | | 517 | 0.19 | 0.23 | 73 | 0.01 | 0.17 |
| LMTV5 - LIPS | | 517 | 0.16 | 0.22 | 73 | 0.15 | 0.18 |
| LIPS - LFEF | | 517 | 0.27 | 0.24 | 73 | 0.11 | 0.19 |
